# Trait-specific responses to repeated stressor exposure in wild crickets

**DOI:** 10.1101/2025.07.14.664656

**Authors:** Emily Gilford, Ruonan Li, Rolando Rodríguez-Muñoz, Tom Tregenza, Bram Kuijper

## Abstract

Habituation allows animals to reduce their responsiveness to repeated, non-threatening stimuli. While well-studied in vertebrates, little is known about how wild insects adjust their behaviour to repeated stressors in natural settings. This study tested whether wild field crickets (*Gryllus campestris*) habituate or sensitise to repeated simulated predator attacks, and whether these effects vary with threat intensity. Crickets showed limited evidence of habituation. Emergence time increased slightly over repeated exposures, consistent with modest sensitisation, while post-emergence distance also increased, suggesting a modest habituation effect. Escape speed remained stable. Our results suggest trait-specific behavioural plasticity, as only some response measures changed across repeated exposures while others remained stable. Crickets ranged further from the burrow in response to the stronger stimulus, suggesting greater comfort or reduced perceived risk, though emergence latency and escape speed were unaffected by stimulus intensity. Contrary to our predictions, individual crickets did not respond consistently across the two stimulus types. While some increased their emergence time or post-emergence distance more under one stimulus than the other, others showed little or no change. These patterns suggest that only certain behaviours were flexible, and that individuals varied in how strongly they adjusted these behaviours. Escape speed, by contrast, remained stable. Together, these results indicate limited habituation, trait-specific flexibility, and substantial individual variation in how crickets respond to repeated disturbance, highlighting the diverse strategies animals may use to manage risk in unpredictable environments.

## Introduction

Natural environments are inherently dynamic, exposing animals to unpredictably fluctuating stressors, from abiotic factors like temperature extremes to biotic threats such as predation and competition (Wingfield and Kitaysky, 2002). Detecting and appropriately responding to these fluctuations is crucial for survival, driving the evolution of diverse adaptive mechanisms (Scheiner, 2013). Responding rapidly to stressors helps animals avoid risky situations, yet over-reacting to stressors may similarly result in opportunity costs and the accumulation of damage (Blumstein, 2016). One mechanism that allows animals to adjust their responses to recurrent stressors is habituation, a form of behavioural plasticity in which repeated exposure to a non-threatening stimulus leads to reduced behavioural responses (Thompson and Spencer, 1966a). Habituation can be adaptive, as stress responses carry well-documented costs, including suppressed immune function, reduced reproduction, and impaired growth (Adamo, 2012; Bonier et al., 2009; Dhabhar, 2014); failure to modulate responses to benign cues could reduce fitness. Yet habituating too quickly to genuine threats may increase vulnerability. Understanding how invertebrates navigate this trade-off is essential for grasping the generality and evolution of stress-coping strategies. By habituating to benign, recurring stimuli, animals conserve valuable energy and physiological resources, reallocating them to fitness-enhancing processes such as growth, reproduction, and immunity (Blumstein, 2016; Rankin et al., 2009).

Studies of habituation have largely focused on vertebrates, finding evidence of habituation in a broad range of taxa, reflecting its fundamental evolutionary significance in managing frequently recurring, low-risk stimuli. The presence of habituation has received particular attention in the context of human disturbance, with many vertebrates, including mammals and birds, readily habituating to non-threatening human presence, conserving energy and reducing unnecessary vigilance. Elk (*Cervus canadensis*) reduce their flight initiation distance with repeated, harmless encounters with hikers, while urban pigeons (*Columba livia*) tolerate close human approach (Carrete and Tella, 2011). Reptiles also show reduced escape responses following repeated non-lethal disturbance (Samia et al., 2016), and harbour seals (*Phoca vitulina*) become progressively less responsive to repeated boat traffic, mitigating energetic costs (Andersen et al., 2012). Beyond human disturbance, habituation to ecologically relevant threats has also been documented: lizards exposed to low-risk predators reduce their flight initiation distance (Rodrı guez-Prieto et al., 2010), and fathead minnows (*Pimephales promelas*) display reduced antipredator behaviour after repeated exposure to weak chemical cues (Dupuch et al., 2004; Zhao and Chivers, 2005). These examples underscore habituation as a broadly conserved strategy, enabling animals to reduce costly responses to low-risk cues and thrive in environments with frequent, non-lethal disturbance.

Although habituation has been documented across a wide range of vertebrate taxa, research on invertebrates, despite their ecological dominance and diversity, has focused primarily on laboratory studies using model species. Taxa such as *Aplysia* (Glanzman, 2009), *Caenorhabditis elegans* (Rose and Rankin, 2001), and *Drosophila* (Trisal et al., 2022) have yielded valuable insights into the neural and mechanistic basis of habituation. However, these studies typically involve restrained or isolated animals exposed to simplified stimuli in tightly controlled environments, which may not reflect the complexity of real-world conditions. Much less is known about how invertebrates adjust their behaviour to repeated stressors in natural contexts. The natural environment introduces variation in context, background risk, and individual experience, factors known to shape behaviour but difficult to replicate in the lab. Without field-based studies, we risk overlooking how invertebrates manage risk in dynamic, ecologically realistic settings.

To our knowledge, only three studies on grasshoppers have directly examined how wild insects respond to repeated predator-associated stressors in naturalistic settings. The first found no evidence of habituation in *D. carolina*, with individuals maintaining strong escape responses regardless of repeated exposure (Cooper Jr., 2006). The second showed context-dependent changes in escape tactics in *P. atlantica*, influenced by prior self-amputation of a limb (Bateman and Fleming, 2011). A third study, again on *P. atlantica*, reported contrasting responses in two grasshopper species: one became less responsive over time, while the other became more evasive, suggesting both habituation and sensitisation can occur, depending on species and context (Bateman and Fleming, 2014). Notably, even these studies were conducted under semi-natural or controlled field conditions, where predator encounters, habitat structure, and individual movement were still partially constrained. These rare examples highlight how little we know about behavioural plasticity in truly wild, free-living invertebrates. This is a critical gap, especially given that many invertebrates are highly sensitive to environmental fluctuations (Stillman, 2003) and rely on rapid behavioural adjustments for survival (Hoffmann and Bridle, 2022). Their stress physiology differs markedly from vertebrates, raising the question of whether patterns of habituation observed in vertebrates, such as reduced responsiveness to repeated, low-risk stimuli, generalise to invertebrates, which is what we set out to assess here. Studying these responses under natural conditions is therefore essential, not only for understanding the functional relevance of behavioural plasticity, but also for predicting how invertebrates may cope with the dynamic and unpredictable environments they face.

Classic models of habituation propose that responses to stressor cues decline monotonically with repeated exposure, particularly when cues are low in salience or risk (Rankin et al., 2009; Thompson and Spencer, 1966a). In contrast, more intense or biologically meaningful cues may elicit sensitisation, where behavioural responses become more pronounced over time. While this divergence is well recognised in theory, empirical studies testing how stimulus intensity shapes the direction of plasticity remain limited, especially in non-model organisms and natural contexts. Another key question concerns the specificity of these responses. Habituation is often assumed to generalise across stimuli, such that repeated exposure to one predator cue reduces responses to similar future threats (Cyr and Romero, 2009; Harnad, 1987). However, generalisation is not universal. In some systems, behavioural changes are highly cue-specific, with responses diminishing only to the repeated stimulus. For example, rodents habituated to one predator odour retain strong responses to odours from other predators (Dielenberg and McGregor, 2001, 1999), suggesting that habituation can be finely tuned to threat identity. Comparable specificity has been found in *Caenorhabditis elegans*, which show site-specific habituation to tactile stimuli (Rose and Rankin, 2001). Yet most invertebrate studies have focused on tactile input, with very limited work investigating generalisation across vibrational or other mechanosensory cues. As such, we still lack a clear understanding of how stimulus intensity, modality, and identity shape the trajectory of behavioural responses to repeated stressors, whether through habituation, sensitisation, or more complex combinations of the two. Addressing this gap will help clarify the functional significance of behavioural flexibility in dynamic and information-rich environments.

This study addresses two key gaps in the habituation literature: first, the scarcity of field-based investigations into how wild invertebrates adjust behaviour over repeated stressor exposures; and second, the limited understanding of individual variation in habituation and sensitisation, which we address by integrating repeatability and plasticity analyses within a multivariate framework.

Field crickets (*G. campestris*) offer an ideal model system for examining how behavioural responses to stressors change over time in natural conditions. As insects that use burrows as a refuge from predation, their antipredator strategy is well-defined: while crickets normally reside outside their burrows to forage or pursue mates, when disturbed, they flee into their burrows. Consequently, this allows for consistent, repeatable measurements of escape behaviour in natural settings (Broder et al., 2024; Stevenson et al., 2000), making crickets particularly well suited to investigating how threat responses change with experience and stimulus intensity.

To this end, we conducted a field experiment to examine how wild crickets in a meadow in northern Spain adjust their responses to repeated disturbances. Crickets were exposed to repeated vibrational stimuli simulating an approaching predator, delivered as either a weaker or stronger cue. We then assessed how crickets respond to these stressor-related cues by measuring key behaviours related to how they flee into their burrows and subsequently re-emerge. We assessed whether crickets showed reductions in escape behaviours over time, consistent with habituation, and whether more intense stimuli elicited stronger or more persistent responses. We focussed on three behaviours which differed in their ecological roles: emergence time, escape speed, and post-emergence distance. Emergence time, the time taken to re-emerge from the burrow post-stressor exposure, reflects caution and environmental risk but also delays access to resources such as food and mates. Escape speed, the speed at which a cricket flees back into the burrow, captures immediate escape intensity with associated energetic costs due to muscular effort. Post-emergence distance, how far the cricket moved away from the burrow entrance after re-emerging, reflects risk perception balanced against the energetic expenditure and potential exposure to further threats that come with movement away from the burrow. Crucially, we were also interested in whether individuals varied in their response trajectories, for instance whether some became less responsive over time (habituation), while others became more responsive (sensitisation). Such differences could reflect variation in behavioural flexibility or inconsistent coping strategies that animals use to manage risk.

As the stressor-related cues used in this experiment posed no actual harm, and therefore represented a low-risk stressor, we predicted that crickets would reduce their responses over time, indicating habituation. In exploring the scope for plastic responses to cues of differing intensity, we predicted that crickets exposed to stronger stimuli would exhibit more pronounced initial responses, longer emergence latencies, faster escape speeds, and greater post-emergence distance from the burrow, compared to those exposed to weaker stimuli. We also expected crickets exposed to weaker stimuli to habituate more rapidly, showing a decline in behavioural responses across repeated trials. Finally, drawing on stimulus generalisation theory (Thompson and Spencer, 1966b), we anticipated consistent individual-level variation in response trajectories, reflecting stable differences in how animals regulate behavioural investment in response to perceived threat.

## Methodology

### Study system

Our study site is the “Wild Crickets” meadow in northern Spain, in which a wild population of the field cricket *G. campestris* has been observed since 2006 (see www.wildcrickets.org) (Tregenza et al., 2022). *G. campestris* are flightless and spend much of their lives around burrows, which they excavate as nymphs (Vrenozi and Uchman, 2020). These burrows provide shelter throughout development, including during winter diapause. Nymphs re-emerge from their burrows in spring and gradually become more active as they grow, reaching adulthood in April and May (Rodríguez-Muñoz et al., 2011). While adult crickets move frequently between burrows to mate (Tregenza et al., 2022) and occasionally dig new burrows, nymphs remain largely restricted to their overwintering burrows, typically foraging within ten centimetres of the entrance (pers. obs.). To regulate their body temperature, crickets bask outside in order to heat up in the sun, and retreat underground during colder or rainy periods or to avoid overheating (Gardner et al., 2024). *G. campestris* are sensitive to substrate-borne vibration (Niemelä et al., 2015), and a recent experiment by Li et al (Li et al., 2025b) at the same study site demonstrated that vibrational stimuli reliably triggered rapid flight into the burrow, confirming this behaviour as an anti-predator response. Using high-speed video and a dropped ball to generate air and vibrational cues, they found that escape speed varied with body temperature in females but not males, further validating escape speed as a reliable, ecologically meaningful response metric.

We conducted field experiments in spring 2024, using *G. campestris* nymphs already present in the meadow from the 24th of March to the 24th of April. In March, we searched the meadow for cricket burrows, which were identified with a uniquely numbered flag, and both burrows and walking paths were mapped. We calculated the distance between each burrow and its nearest path in metres using spatial data processed in QGIS. This “path distance” variable was included in subsequent analyses to examine whether burrow location, and by extension, potential prior exposure to disturbance, affected crickets’ behavioural responses to repeated stressor-related cues. We installed infra-red cameras over 140 of the burrows, monitoring activity 24 hours a day, allowing behaviours and developmental stages of crickets to be tracked.

### Cues of predation and response measures

We followed a similar methodology to that described in Li *et al*, in which a 240 frames per second, 2.7K pixel GoPro was used to produce detailed recordings of fleeing events (Li et al., 2025b). We mounted the camera on a tripod, 50 cm above the substrate and facing directly downwards above the burrow. We placed a 30 cm ruler, accurate to the nearest millimetre, adjacent to the burrow to serve as a reference for distance measurements. A 50 cm tube was also affixed to the tripod, with the end held 15 cm above the ground, pointing approximately 10 cm away from the cricket burrow’s entrance. We delivered an artificial air current and vibrational stimulus by dropping a ball down the tube. The experimental setup (Figure S1) standardised the magnitude of vibrations reaching the cricket. To test for changes in flight responses over time in relation to stimulus intensity, we used two types of artificial vibrational stimuli across up to 10 successive trials. The stimuli comprised 42 mm diameter balls: a 9 g cork ball and a 266 g steel ball. These generated vibrational cues of differing intensity, with the cork ball producing a weaker stimulus and the steel ball a stronger one. We collected all data (n = 438 trials) from 37 individual crickets, each identified by a unique burrow. Each individual experienced up to 10 trials per stimulus type, with a total of 214 trials using the weak stimulus, and 224 using the strong. Trials with each ball type were conducted on separate days, allowing at least one day of rest between treatments. The order of stimulus exposure was alternated between individuals to minimise potential order effects.

An operator set up the apparatus and sat quietly near the burrow, waiting for a cricket to emerge. Once the cricket emerged and remained still for 1 minute, the operator measured its body temperature using an infrared thermometer (Testo 830-T4; see https://www.testo.com/). A laser pointer marked its position with a red dot visible to the naked eye before applying the stimulus. We started the high-speed video recording remotely via smartphone trigger and immediately released the stimulus.

We measured three behavioural responses to repeated predator-mimicking stimuli: emergence time, escape speed, and post-emergence distance. Emergence time was recorded as the latency for a cricket to exit its burrow following disturbance, escape speed measured the rapidity of movement as the cricket fled into its burrow, and post-emergence distance quantified how far the cricket moved away from the burrow entrance after re-emerging, and before remaining still for 1 minute. These metrics capture different aspects of the crickets’ antipredator behaviour related to risk assessment and energetic investment. All behaviours were quantified using high-speed video recordings and analysed frame-by-frame to obtain precise timing and distances. Together, these measures provide insight into how crickets modulate defensive behaviour in response to repeated stressor exposure.

We recorded emergence time as the interval between when the cricket entered the burrow and when its entire body emerged. Once the cricket had again remained still outside the burrow for 1 minute, a second trial could take place, up to a maximum of 10 trials per cricket per stimulus intensity. If emergence time exceeded 10 minutes, or if the cricket took longer than 10 minutes to remain still enough for the next trial to take place, then the iteration would end to minimise chronic stress.

### Video Analysis

We extracted distance fled and escape speed from video recordings using ‘Kinovea’ (https://www.kinovea.org/download.html), which allows frame-by-frame analysis, with each frame representing 1/240^th^ of a second.

We measured post-emergence distance in centimetres, which reflected the furthest point the cricket reached from the burrow entrance after re-emerging in response to the stressor-related cue, up until it became still for 1 minute. We calculated escape speed over the first 1.5 cm of movement. All individuals ran for at least 1.5cm, so standardising to this distance avoids the effects of speed increasing with distance. Our approach follows a previous experiment at the same field site (Li et al., 2025b), which used a similar vibrational stimulus and found 1.5□cm to be sufficient for capturing early escape behaviour, while minimising variation due to turning angle, distance to refuge, or variation in initial distance from the burrow. We extracted the following parameters from the video: (1) the video frame in which the cricket’s first obvious movement was detected, which we denote as the response frame (*rf*), (2) the video frame in which the stressor-related cue was released (release frame), (3) the frame in which the cricket reached a distance of 1.5cm relative to the position of the cricket when the stressor-related cue was released (*t_1_*) and (4) the frame that contained the maximum distance fled. Parameters were used to calculate escape speed (m/s). We calculated the time taken for a cricket to cover 1.5cm (*t_x_)* using *t_x_* = (*t_1_ − rf*)*(1/240)). We then calculated escape speed across the 1.5cm segment to calculate the escape speed in meters per second (*fs*); fs = 0.015 / *t_x_*.

### Data Analysis

To examine response changes across trials, we included only burrows with at least four trials for either stimulus type. We chose this threshold to ensure sufficient repeated measures for estimating individual behavioural variation with reasonable accuracy and statistical power; previous work shows that at least 4–5 repeats are typically required to obtain reliable estimates of within-individual variance and behavioural repeatability (Dingemanse and Dochtermann, 2013). The resulting dataset comprised 37 cricket nymphs (17 females, 19 males, 1 unknown), totalling 438 observations. Of these, 11 burrows were exposed only to the weaker stimuli, 11 only to the stronger stimuli, and 16 to both. We included individual ID as a random effect in all analyses to avoid pseudoreplication and to estimate individual-level response changes, as captured by a random slopes effect.

We fitted Bayesian multivariate mixed-effects models in R (v4.3.2)(R Core Team, 2023) using the *brms* package (Bürkner, 2017), with emergence time, escape speed, and post-emergence distance as response variables. All variables were continuous and right-skewed, but Box-Cox analyses (MASS package (Venables and Ripley, 2002) indicated minimal transformation was required (λ ≈ 1). Simplified models were first compared to select the most appropriate GLM family, with posterior predictive checks confirming superior predictive accuracy of Gaussian models relative to Gamma or log-normal alternatives. Although residual normality is often emphasised, recent evidence suggests that Bayesian mixed-effects models are robust to slight violations (Gelman and Hill, 2006; McElreath, 2020; Schielzeth et al., 2020), supporting the use of a Gaussian likelihood in this context. Posterior predictive checks showed close alignment between observed and simulated data for all three response variables, with no major deviations from normality (Figure S2). Residuals were symmetrically distributed and exhibited no obvious patterns of heteroscedasticity or bias, further justifying the Gaussian specification.

We centred response variables within each burrow and stimulus combination (by subtracting the mean per burrow–stimulus combination) to separate within-burrow variation (e.g., across trials) and between-burrow variation (Dingemanse and Dochtermann, 2013; van de Pol and Wright, 2009). Additionally, we multiplied post-emergence distance by -1 (sign reversal) to reflect that smaller distances indicate habituation, aligning it with the other two response variables. Therefore, negative model estimates correspond to increases in the raw distance crickets travelled after re-emerging. Trial number, stimulus type, temperature, sex, path distance, and an interaction between trial number and stimulus type were included as predictors to capture how behavioural responses changed across trials depending on stimulus intensity. All ordinal (trial) and continuous (temperature, path distance) independent variables were mean-centred, and we effect-coded stimulus intensity as ±0.5 (strong = 0.5, weak = – 0.5). Sex was included as a three-level factor (female, male, unknown), with “female” as the reference level. This approach improves interpretability of main effects in the presence of interactions while retaining all individuals in the analysis (Kraemer and Blasey, 2004; Schielzeth, 2010). Random slopes for trial, stimulus type, and their interaction were specified within burrow using the structure (trial * stimulus | p | burrow), allowing separate variance-covariance matrices for each response and enhancing model flexibility (Bürkner, 2018).

### Ethics

This study was approved by the University of Exeter Research Ethics Panel (approval number: 3941951). A total of 37 cricket nymphs of both sexes were used. Each experimental session lasted no more than one hour, with a minimum rest period of one day between stimulus types. Trials were terminated immediately if crickets did not re-emerge after ten minutes, attempted to flee, or failed to remain still, to minimise chronic stress. Following the study, all crickets were released to live out their natural lives in the wild. No crickets were harmed during the experiment.

## Results

### Do crickets adjust their behavioural responses to stressors across successive trials?

We found that crickets only slightly modulate their responses during repeated exposure to the same stressor-related cue (Figures 1 and 2; Table S1). Crickets showed a small but credible increase in emergence time across repeated trials (β_trial_ = 2.48, 95% CI [0.21, 4.77]; Table S1), consistent with a modest habituation effect, in which individuals become less cautious over time. Crickets did not differ in their escape speed over successive trials (β_trial_ = −0.00, 95% CI [−0.01, 0.00]; Figure 2B, Table S1), suggesting that the speed at which they fled is constant and neither subject to habituation nor sensitisation. Finally, crickets slightly decreased their post-emergence distance over successive trials (β_trial_ = 0.06, 95% CI [0.03, 0.09]; Figure 2C, Table S1), consistent with a modest but credible sensitisation effect. The field experiment also considered other variables influencing responses to stressor-related cues, including temperature, sex, and path distance. However, credible intervals for these predictors consistently overlapped zero across all traits (Figure 2, Table S1), indicating no detectable effect on behavioural responses.

**Figure 1.**
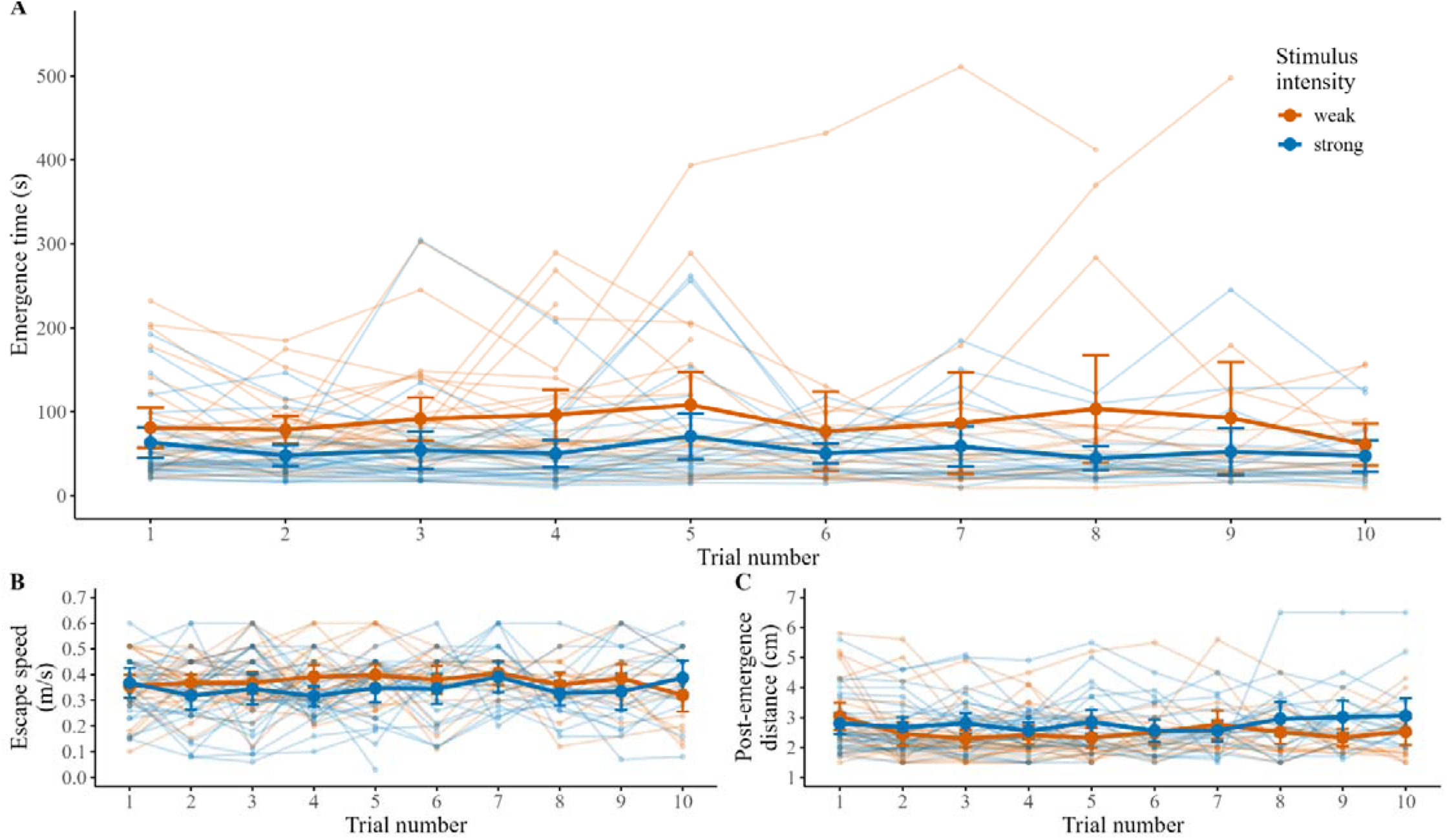
Behavioural responses across repeated trials. Line graphs showing (A) emergence time (s), (B) escape speed (m/s), and (C) post-emergence distance from burrow (cm) for individuals across 10 repeated trials. Thin lines represent individual trajectories, while bold lines show mean values ± 95% confidence intervals for each trial. Colours indicate stimulus intensity (orange: weak; blue: strong).

**Figure 2.**
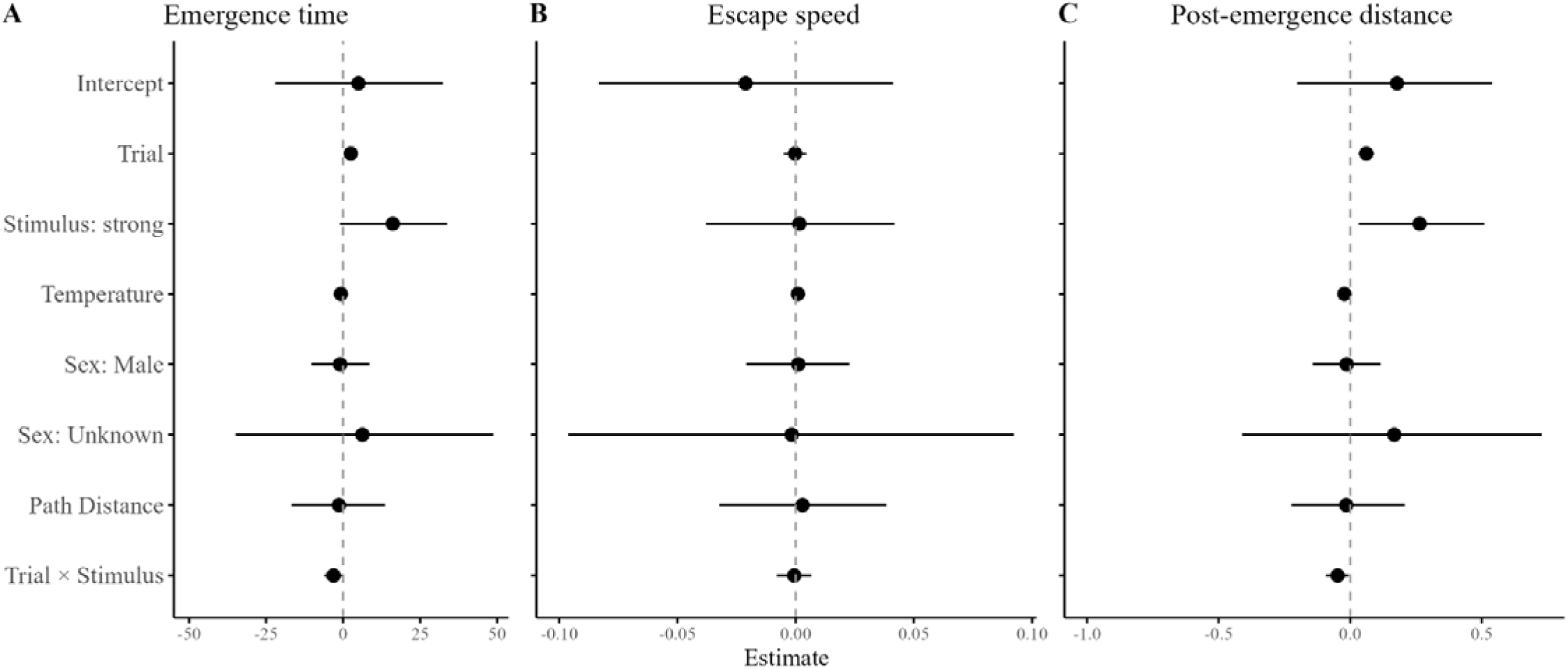
A forest plot showing fixed effect estimates for emergence time, escape speed, and post-emergence distance. Posterior mean estimates (black dots) and 95% credible intervals (black horizontal lines) for the model parameters: trial, stimulus type (strong), path distance, sex (male), and temperature for (A) emergence time. (B) escape speed and (C) post-emergence distance. The vertical dashed line at zero indicates no effect for each parameter.

### Response variation depending on stressor intensity

Stimulus intensity also influenced cricket responses across trials (Figure 1, Table S1). When examining stimulus intensity alone, crickets tended to show higher emergence times in response to stronger stimuli relative to weaker ones (see Figure 2: β_stimulus:strong_ = 16.08, 95% CI [−1.56, 34.12]; Table S1), though this effect was highly uncertain and not credibly different from zero. Similarly, crickets exhibited slightly faster escape speeds when exposed to the stronger stimulus compared to the weaker one, though the magnitude of the difference was small and again not credibly different from zero (β_stimulus:strong_= 0.00, 95% CI [−0.04, 0.04]; Table S1). Post-emergence distance was credibly affected by stimulus intensity: individuals exposed to the stronger stimulus settled closer to the burrow following emergence than those exposed to the weaker stimulus (β_stimulus:strong_= 0.26, 95% CI [0.02, 0.51]; Table S1). A significant trial-by-stimulus interaction (β_trial:stimulus_ = −0.05, 95% CI [−0.09, −0.00]; Table S1) revealed divergent patterns over time, while crickets under the weaker stimulus gradually reduced their post-emergence distance consistent with a modest habituation response, those exposed to the stronger stimulus maintained or increased this distance, suggesting slight sensitisation. Emergence time also showed a credible interaction between trial number and stimulus intensity, with crickets exposed to the stronger stimulus tending to delay emergence increasingly over repeated exposures, again consistent with sensitisation to more intense stressor conditions (β_trial:stimulus_= −3.09, 95% CI [−6.30, −0.01]; Table S1).

### Plasticity in responses to stressor intensity

We then assessed whether individuals differed in their degree of habituation or sensitisation when exposed to different stimulus intensities. To do this, we compared individual-level slopes of behavioural responses across trials for crickets exposed to both weak and strong stimuli in successive experiments (n = 17; Figure 3A–C). For all three behaviours, emergence time (Figure 3A), escape speed (Figure 3B), and post-emergence distance (Figure 3C), slopes under weak and strong stimulus conditions were only weakly correlated (emergence time: r = −0.02; escape speed: r = −0.04; post-emergence distance: r = 0.03), suggesting little consistency in individual plasticity across contexts. Paired t-tests showed no significant differences in mean slope between weak and strong stimuli for any behavioural metric (emergence time: *t*□□ = 1.32, *p* = 0.207; escape speed: *t*□□ = −0.40, *p* = 0.693; post-emergence distance: *t*□□ = 1.05, *p* = 0.310), indicating that stimulus intensity did not systematically alter the degree to which individuals habituated or sensitised across repeated trials.

**Figure 3.**
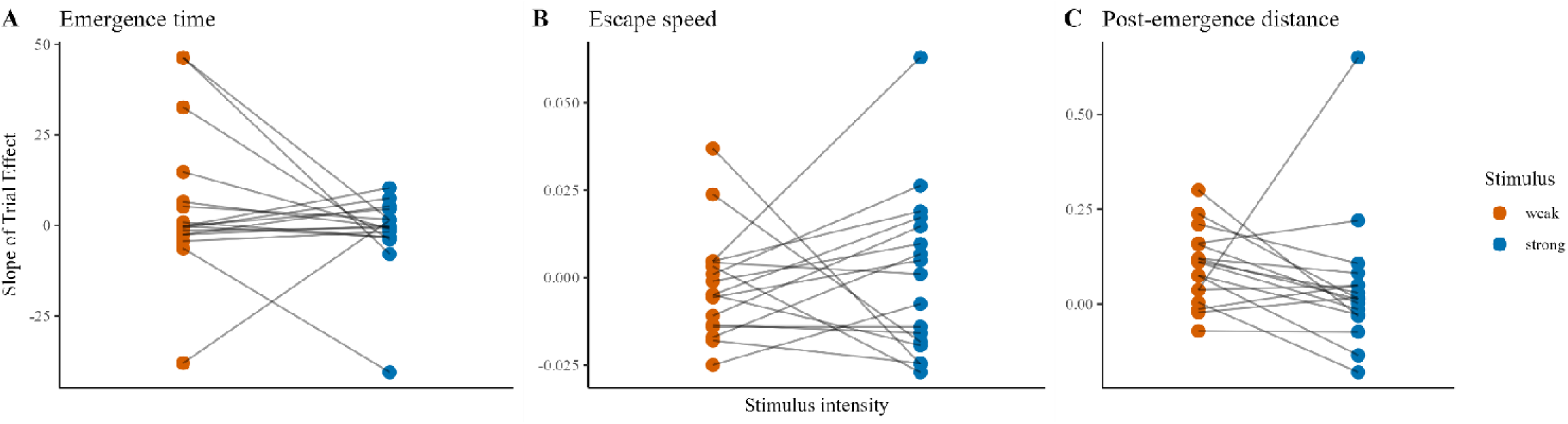
Random slope estimates for behavioural responses across stimulus strengths. Individual slope estimates for (A) emergence time, (B) escape speed and (C) post-emergence distance, comparing weak (orange) and strong (blue) stressor intensities. Points represent individual slopes, with lines connecting paired data points for stimulus intensity types. Includes individuals who experienced both stimulus intensities (n = 17).

### Between-individual variation in response change

We found substantial between-individual variation in behavioural responses across trials, as indicated by the standard deviations of the random slopes (Table S2). For emergence time, there was no clear main effect of stimulus intensity across individuals (β = 16.08, 95% CI [−1.56, 34.12]; Table S1)

However, the interaction between trial and stimulus was credibly different from zero (β = −3.09, 95% CI [−6.30, −0.01]; Table S1), indicating that the direction or magnitude of change in emergence time across trials differed depending on stimulus strength. Specifically, emergence time increased more over repeated exposures to the weak stimulus, consistent with modest habituation, while this pattern was less pronounced, or possibly reversed, under the strong stimulus. Individual responses were highly variable, with random slope standard deviations of 0.70 for trial, 3.09 for stimulus intensity, and 0.63 for their interaction (Table S2), reflecting considerable heterogeneity in behavioural plasticity across individuals (Figure 2).

A similar interaction pattern was observed for post-emergence distance. Crickets exposed to the stronger stimulus showed greater distances from the burrow across trials (β_trial:stimulus_ = −0.05, 95% CI [−0.09, −0.00]; Table S1), consistent with a modest habituation effect under strong stimulation. Random slope standard deviations were 0.01 for both the trial and the trial-by-stimulus interaction terms (Table S2), indicating a small but present level of between-individual variation in plasticity.

In contrast, escape speed remained stable across trials and stimulus intensities. Fixed effect estimates for trial (β = 0.00, 95% CI [−0.01, 0.00]) and for the interaction between trial and stimulus (β = 0.00, 95% CI [−0.01, 0.01]; Table S1) were both close to, and not credibly different from, zero. Corresponding random slope standard deviations were also minimal (0.00–0.01 across terms; Table S2), suggesting that this behavioural response was largely consistent across individuals and repeated exposures.

## Discussion

### Behavioural change across repeated exposures

We sought to assess whether crickets showed reductions in escape behaviour over time, consistent with habituation. Across repeated exposures to either weaker or stronger stressor-related cues, crickets did not exhibit uniform habituation in their escape behaviours, and no single population-level pattern emerged. Instead, we observed slight but credible increases in both emergence latency and post-emergence distance over trials, indicating that crickets took longer to emerge from their burrows and travelled further once they did, respectively. The former pattern suggests a modest sensitisation effect, reflecting increased caution over time, while the latter suggests modest habituation, as individuals ranged further from the burrow following repeated exposures. Escape speed, by contrast, remained stable, showing no consistent directional change at the population level.

These findings diverge from our original prediction that crickets would habituate to repeated stimuli, showing progressively reduced behavioural responses over successive trials. Instead, the direction and magnitude of behavioural change appeared to vary across traits and individuals. This finding runs counter to classical models of habituation, which predict a monotonic decrease in response strength with repeated exposure to a non-threatening stressor-related cue (Rankin et al., 2009; Thompson and Spencer, 1966a). Moreover, rather than reflecting a single population-level trajectory, our data reveal substantial variation in how individuals responded over time. This echoes findings from studies in both vertebrates and invertebrates that exhibit varied stress-related plasticity (Adamo, 2012; Cyr and Romero, 2009; Zhao and Chivers, 2005). The relative stability of escape speed, compared to the increases in emergence time and post-emergence distance, suggests these traits may serve different ecological roles: for instance, escape speed may reflect a more hardwired motor response shaped by biochemical limits, while emergence time and post-emergence distance may be more flexible and context-dependent, reflecting environmental appraisal, perceived risk, and investment in further movement once threats subside (Biro and Stamps, 2008).

Sensitisation may reflect a behavioural response to an accumulation of physiological or psychological stress over repeated exposures, particularly in a natural setting where misclassifying threats can have high costs (Blanchard et al., 1995; Helfman, 1989). Unlike in tightly controlled laboratory conditions, repeated vibrational cues in the field may not be easily categorised as benign, leading to persistent uncertainty or heightened vigilance (Brown et al., 2011). Additionally, the functional roles of these behaviours may differ: emergence time likely reflects a decision-making process influenced by continued perception of environmental risk and internal state (Lima and Dill, 1990), whereas escape speed may be a more instantaneous and hence more reflexive response (Domenici and Blake, 1997). These differences mirror broader patterns seen in prey species, where more deliberative behaviours are often more plastic and context-sensitive than rapid escape responses (Blumstein, 2016). The slight sensitisation of emergence time, contrasted with the modest habituation of post-emergence distance and the stability of escape speed, highlights that different components of antipredator behaviour may respond differently to repeated disturbance, and that only some traits may be flexibly regulated in response to threat perception.

### Threat intensity and behavioural modulation

We predicted that crickets exposed to the stronger stimulus would show stronger initial responses such as longer emergence latencies and faster escape speeds, alongside reduced post-emergence distance, remaining closer to the burrow due to heightened risk perception. Contrary to this, crickets tended to show higher emergence times in response to the stronger stimulus (Figure 2A), though this effect was highly uncertain and not credibly different from zero. Similarly, crickets exposed to the stronger stimulus showed slightly faster escape speeds (Figure 2B), but again, this difference was small and not credible. However, crickets exposed to the stronger stimulus settled closer to their burrows following emergence than those exposed to the weaker stimulus (Figure 2C), and this effect was credibly different from zero. We observed a credible interaction between trial and stimulus intensity, revealing that while crickets under the weaker stimulus reduced their post-emergence distance over time, consistent with a modest sensitisation response, those exposed to the stronger stimulus showed increasing post-emergence distances, suggesting slight habituation. A similar pattern was observed for emergence time, which also showed a credible interaction effect, with crickets delaying emergence more across trials when exposed to stronger stimuli.

These results suggest that crickets do not respond to vibrational cues in a strictly intensity-dependent way. One possibility is that the weaker cork ball stimulus resembles biologically relevant predator cues, such as subtle vibrations from insectivorous birds like robins, while the stronger steel ball stimulus may be perceived as a non-predatory disturbance (e.g. large mammals or human footfall), prompting a less urgent initial response. This interpretation aligns with the threat-sensitivity hypothesis (Helfman, 1989) and the predation risk allocation hypothesis (Lima and Bednekoff, 1999), both of which emphasise the role of risk interpretation over raw cue intensity. Our findings also partially mirror previous studies in *G. campestris*, which observed slower escape speeds in response to stronger vibrational cues (Li et al., 2025a, manuscript submitted for publication). Although the effect size reported in that study was small (−0.044□m/s), it was statistically detectable. In contrast, our estimate (β_stimulus:strong_ = 0.00; 95% CI [−0.04, 0.04]) shows no credible difference, which may reflect differences in sample size, statistical power, or study design. This is also consistent with the narrow range and relative stability of escape speed across trials observed in our data. Together, these results suggest that crickets modulate behaviour according to the perceived meaning of a threat rather than its intensity alone. In contrast to our prediction, individuals exposed to both intensities of stressor-related cue did not respond in consistent ways (Figure 3), with weak and inconsistent correlations across all three traits. This suggests responses did not generalise from one cue to another. Correlations across traits were weak and inconsistent, providing little evidence that individual behavioural responses to vibrational stimuli were stable across different cue types. As such, we cannot determine whether these responses reflect cue-specific plasticity or broader generalisation processes.

### Variation in individual plasticity under repeated threat occurrence

In addition to variation across stimuli and over time, we found individual-level differences in behavioural trajectories (Figure 1), supporting our prediction that individuals would differ consistently in how they adjust their responses across repeated exposures. Some crickets habituated, showing reduced responsiveness over time, while others exhibited patterns consistent with sensitisation. These divergent patterns were reflected in the random slope estimates for emergence time, where credible variation across individuals suggests that behavioural plasticity differs meaningfully among crickets (Figure 3, Table S2). For escape speed and post-emergence distance, random slope estimates were near zero and not credibly different from zero, suggesting little to no evidence of individual differences in plasticity for these traits.

### Trait-specific and variable behavioural plasticity

Our findings show that behavioural plasticity in crickets is trait-specific and individually variable, with some responses, like emergence time and post-emergence distance, showing modest flexibility, while others, such as escape speed, remain more stable. This supports the view that plasticity is multifaceted, likely reflecting differences in the costs, constraints, and ecological roles of different behaviours. Such flexibility may be adaptive in dynamic environments, where adjusting responses appropriately can influence survival and energy use. In our study, increased emergence latency over time in response to stronger stimuli suggests trait-specific sensitisation, while increased post-emergence distance indicates modest habituation. The absence of consistent change in escape speed highlights its relative stability. These patterns highlight the importance of understanding how stressor-related cue characteristics and trait function interact to shape behavioural outcomes.

### Study limitations and directions for future research

One limitation of our approach is the potential for self-selection bias during the initial selection of experimental individuals. We only included crickets that emerged from their burrows within ten minutes before the first trial, which may have excluded individuals with more cautious or neophobic behavioural profiles. As a result, our sample may be biased toward bolder or more disturbance-tolerant individuals. This could lead to an underrepresentation of crickets with low behavioural plasticity or greater sensitivity to environmental cues. Future studies should address this sampling bias by extending observation windows, using automated monitoring, or treating emergence latency as an initial trait of interest.

## Conclusion

Our findings demonstrate that habituation to repeated vibrational stimuli in crickets is trait-specific and context-dependent. Emergence time and post-emergence distance both increased across trials, indicating greater caution and behavioural investment consistent with modest sensitisation for emergence time, and modest habituation for post-emergence distance, whereas escape speed remained stable, suggesting it may be a more reflexive and less plastic trait. These results indicate that behavioural plasticity is shaped by the functional role of each trait rather than occurring uniformly. Additionally, while stimulus intensity did not credibly affect escape speed or emergence latency, it did influence post-emergence distance, with stronger cues prompting crickets to settle closer to the burrow and a trend toward decreasing distances over time. This suggests that crickets may interpret different vibrational cues in distinct ways. By using a field-based approach, our study complements laboratory work and highlights how behavioural plasticity unfolds in more ecologically relevant, variable environments.

