## Supplementary Materials for "Trait-specific responses to repeated stressor exposure in wild crickets"

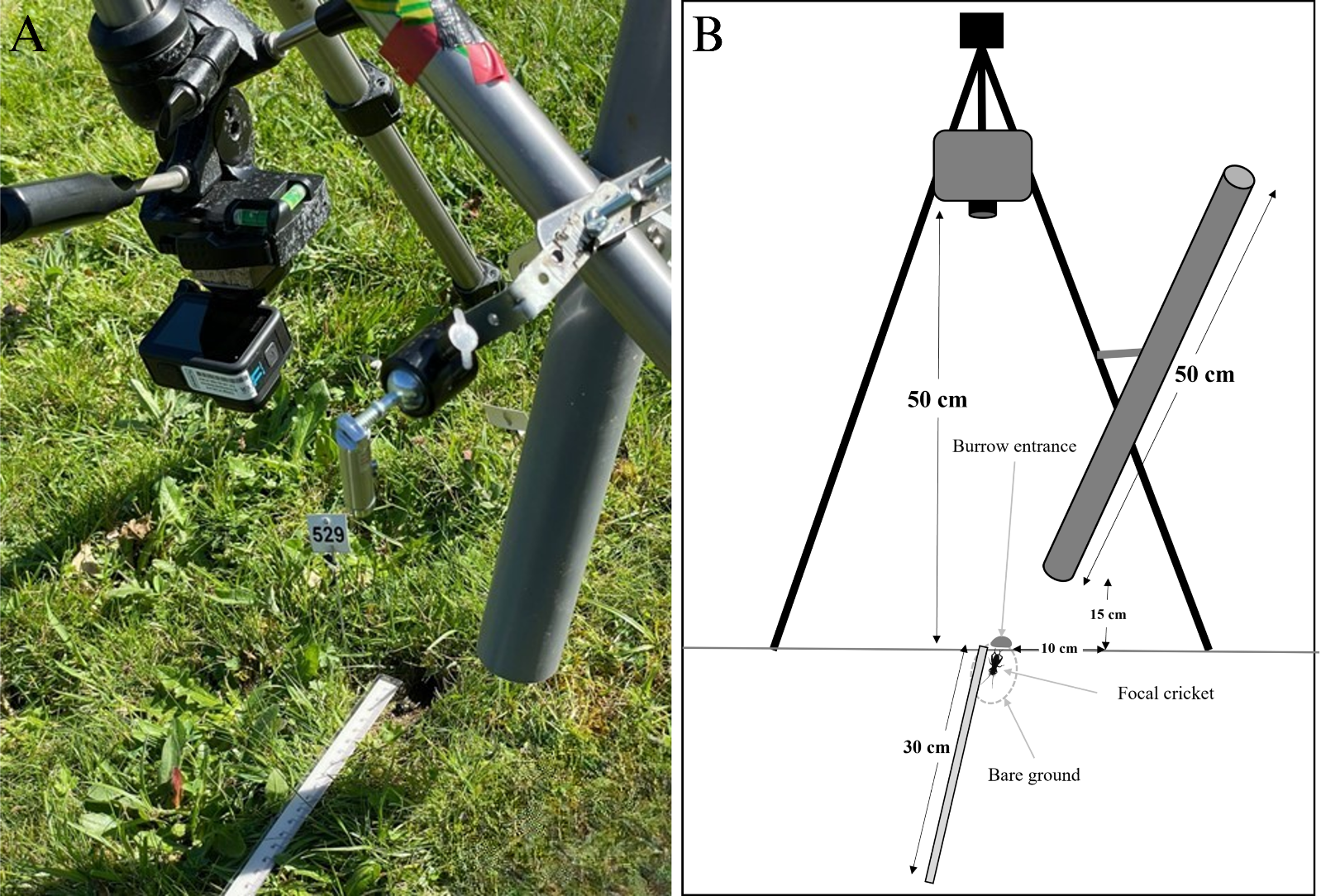

### Methodology

Figure S1. Experimental setup. (A) A tripod-mounted GoPro (240 fps, 2.7K) was positioned 50 cm above an occupied cricket burrow. A 30 cm ruler was placed beside the burrow, aligned with the main path of cricket emergence along a patch of bare ground to allow for distance measurements. The ruler was angled to follow this natural emergence path rather than being parallel to the horizon. (B) Schematic showing the camera setup, ruler placement, and stimulus tube, which ended 15 cm above the ground and delivered vibrational stimuli approximately 10 cm from the burrow entrance. The location of the bare ground and burrow entrance is indicated by a dashed circle**.**.

#### Data Analysis

Analysis was conducted in R (v4.3.2)(R Core Team, 2023). Data and an analysis script can be provided on request. The script performs data cleaning and centring, fits a multivariate Bayesian mixed-effects model (using brms) to assess behavioural responses—emergence time, escape speed, and post-emergence distance from the burrow—to repeated stimuli, and generates posterior predictive plots. It also extracts individual-level plasticity estimates by calculating trial slopes for each stimulus type, and visualises both average trends and individual variation in response. Figures produced include behavioural trajectories, a forest plot of fixed effects, and variation in plasticity across stimuli.

All models were run using four chains with default settings in brms, resulting in 2,000 iterations per chain (1,000 warmup, 1,000 post-warmup), for a total of 4,000 posterior samples. Trace plots for all parameters were inspected to confirm good mixing, and R-hat values were <1.00 across all parameters, indicating convergence. Posterior predictive checks and DHARMa residual diagnostics were also used to assess model fit and revealed no major issues aside from minor deviations in tail behaviour, which were consistent with slight overdispersion rather than model misspecification.

Default priors were used throughout, which was appropriate given the lack of strong prior information in this system. The default priors in brms serve to regularize parameter estimates without overpowering the likelihood, providing a principled balance between flexibility and stability. Posterior distributions showed reasonable behaviour and were not truncated at the bounds of the priors, supporting the adequacy of this approach.

### Results

#### Tables

| **Fixed effects** | *Parameter* | *Estimate* | *SE* | *Lower 95% CI* | *Upper 95% CI* | *Rhat* |
| --- | --- | --- | --- | --- | --- | --- |
| **Emergence time** |  |  |  |  |  |  |
|  | **Intercept** | 4.59 | 14.06 | -23.35 | 32.86 | 1.00 |
|  | Trial | 2.48 | 1.16 | 0.21 | 4.77 | 1.00 |
|  | Stimulus: strong | 16.08 | 9.08 | -1.56 | 34.12 | 1.00 |
|  | Temperature | -0.74 | 0.50 | -1.73 | 0.23 | 1.00 |
|  | Sex: Male | -0.96 | 4.92 | -10.83 | 8.99 | 1.00 |
|  | Sex: Unknown | 5.71 | 20.56 | -34.62 | 46.84 | 1.00 |
|  | Path Distance | -1.37 | 7.84 | -17.06 | 14.45 | 1.00 |
|  | Trial: Stimulus Interaction | -3.09 | 1.60 | -6.30 | -0.01 | 1.00 |
| **Escape speed** |  |  |  |  |  |  |
|  | **Intercept** | -0.02 | 0.03 | -0.08 | 0.04 | 1.00 |
|  | Trial | 0.00 | 0.00 | -0.01 | 0.00 | 1.00 |
|  | Stimulus: strong | 0.00 | 0.02 | -0.04 | 0.04 | 1.00 |
|  | Temperature | 0.00 | 0.00 | 0.00 | 0.00 | 1.00 |
|  | Sex: Male | 0.00 | 0.01 | -0.02 | 0.02 | 1.00 |
|  | Sex: Unknown | 0.00 | 0.05 | -0.10 | 0.09 | 1.00 |
|  | Path Distance | 0.00 | 0.02 | -0.03 | 0.04 | 1.00 |
|  | Trial: Stimulus Interaction | 0.00 | 0.00 | -0.01 | 0.01 | 1.00 |
| **Post-emergence distance** |  |  |  |  |  |  |
|  | **Intercept** | 0.18 | 0.19 | -0.18 | 0.55 | 1.00 |
|  | Trial | 0.06 | 0.02 | 0.03 | 0.09 | 1.00 |
|  | Stimulus: strong | 0.26 | 0.12 | 0.02 | 0.51 | 1.00 |
|  | Temperature | -0.02 | 0.01 | -0.04 | -0.01 | 1.00 |
|  | Sex: Male | -0.02 | 0.07 | -0.14 | 0.12 | 1.00 |
|  | Sex: Unknown | 0.15 | 0.29 | -0.40 | 0.73 | 1.00 |
|  | Path Distance | -0.01 | 0.11 | -0.22 | 0.21 | 1.00 |
|  | Trial: Stimulus Interaction | -0.05 | 0.02 | -0.09 | 0.00 | 1.00 |

Table S1 - Posterior summaries of the fixed effects from the Bayesian mixed-effects model (rsmodel) which tests how the three response variables change over consecutive trials and other variables. The model included emergence time, escape speed, and post-emergence distance as response variables. Predictors were trial number, stimulus intensity, temperature, sex, and path distance. Posterior mean estimates, standard errors, and 95% credible intervals are shown. Rhat values indicate model convergence, with values close to 1.00 suggesting adequate convergence. Each section corresponds to a different behavioural response variable. Intercepts represent the estimated baseline value when all predictors are at their reference levels: trial = 0, weak stimulus, female sex, and mean-centred temperature and path distance.

| **Random effects** | *Parameter* | *Estimate (SD)* | *SE* | *Lower 95% CI* | *Upper 95% CI* |
| --- | --- | --- | --- | --- | --- |
| **Emergence time** |  |  |  |  |  |
|  | **Intercept** | 2.47 | 2.07 | 0.08 | 7.59 |
|  | Trial | 0.70 | 0.54 | 0.03 | 2.03 |
|  | Stimulus: strong | 3.09 | 2.48 | 0.12 | 9.24 |
|  | Trial: Stimulus Interaction | 0.63 | 0.50 | 0.02 | 1.88 |
| **Flee Speed** |  |  |  |  |  |
|  | **Intercept** | 0.01 | 0.00 | 0.00 | 0.02 |
|  | Trial | 0.00 | 0.00 | 0.00 | 0.00 |
|  | Stimulus: strong | 0.01 | 0.01 | 0.00 | 0.02 |
|  | Trial: Stimulus Interaction | 0.00 | 0.00 | 0.00 | 0.00 |
| **Post-emergence distance** |  |  |  |  |  |
|  | **Intercept** | 0.04 | 0.03 | 0.00 | 0.10 |
|  | Trial | 0.01 | 0.01 | 0.00 | 0.02 |
|  | Stimulus: strong | 0.05 | 0.04 | 0.00 | 0.13 |
|  | Trial: Stimulus Interaction | 0.01 | 0.01 | 0.00 | 0.03 |

Table S2 - Posterior summaries of the group-level standard deviations (SDs) for the random effects in the Bayesian mixed-effects model (rsmodel). This model assesses how within and between-individual variation in how the three response variables change over consecutive trials and other variables. The model included emergence time, escape speed, and post-emergence distance as response variables. Random intercepts and random slopes for trial, stimulus intensity (strong), and their interaction were estimated for each response variable, grouped by burrow ID (n = 37). The table reports the posterior mean SDs, standard errors, and 95% credible intervals. These values reflect individual-level variation in behavioural responses across burrows. Intercepts represent baseline behavioural responses when all predictors are at their reference levels: trial = 0, weak stimulus, female sex, and mean-centred temperature and path distance.

#### Posterior Predictive Plots

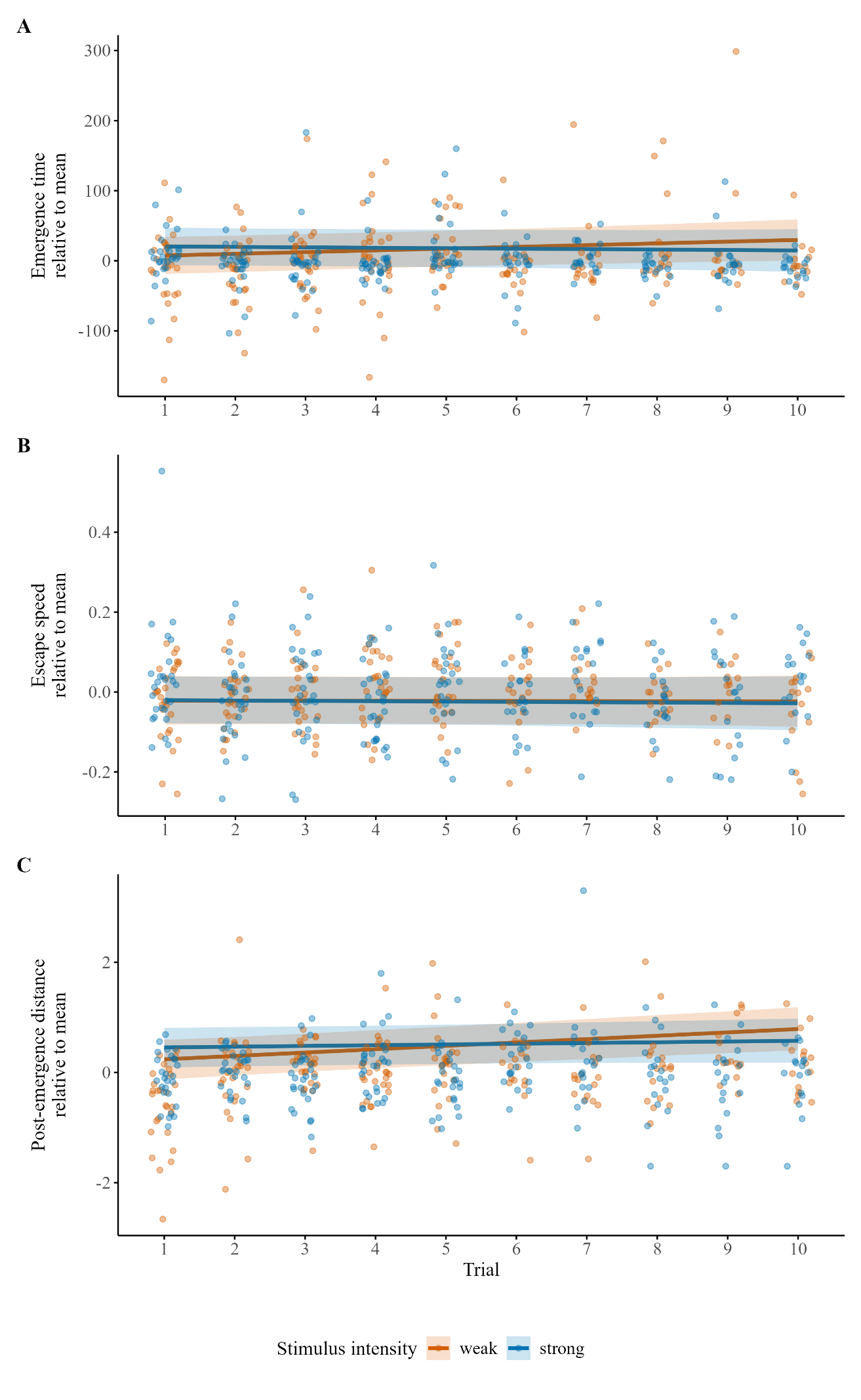

Figure S2 – Posterior predictive plots, showing predicted behavioural responses from the model presented in Tables S1 and S2 across repeated exposures (trial 1–10) to weak (orange) and strong (blue) vibrational stimuli. Solid lines represent posterior mean estimates of the fixed effects; shaded ribbons indicate 95% credible intervals. Jittered points represent raw data for each trial. Plots show: (A) emergence time, (B) escape speed, and (C) post-emergence distance, each centred by their respective means per stimulus type. Data are mean-centred; negative values reflect lower than average values while positive values reflect higher than average values. Predictions were generated with other covariates (temperature, sex, path distance) held constant at reference values (mean or female category) and excluding random effects.

### References

R Core Team, 2023. R: a language and environment for statistical computing (manual). R Foundation for Statistical Computing, Vienna, Austria.
